# Highly Sensitive Chemigenetic FRET-Based Kinase Biosensors

**DOI:** 10.64898/2026.02.06.702961

**Authors:** Xinchang He, Daojia R. Zhou, Yu Huan, Zachary Berriman-Rozen, Sohum Mehta, Yao Chen, Michelle S. Frei, Jin Zhang

## Abstract

Fluorescent protein-based biosensors have transformed the study of cell physiology and pathology by enabling direct, live-cell measurements of biochemical activities with spatiotemporal precision. FRET-based biosensors offer a quantitative and well-defined readout mechanism popular among researchers, but have struggled to break free of characteristically low dynamic ranges and overall dependence on the cyan-yellow spectral region. Chemigenetic approaches that combine synthetic fluorophores with self-labeling protein tags represent an attractive solution to these longstanding constraints. Here, we pair different fluorescent protein donors with a HaloTag acceptor conjugated to a far-red fluorophore to obtain a suite of highly sensitive, chemigenetic FRET-based kinase activity biosensors with red-shifted emission and unprecedented dynamic range. We demonstrate the generalizability of this chemigenetic platform by developing biosensors for multiple kinases, as well as small GTPases and second messengers, all while maintaining high sensitivity. The high sensitivity and spectral tunability of these chemigenetic tools enabled us to perform robust multiplexed activity imaging of receptor-mediated signaling networks to quantitatively map isoform-specific coupling by GPCRs, as well as clear visualization of kinase activity in acute brain slices via two-photon fluorescence lifetime imaging. Our chemigenetic sensor toolkit thus provides the sensitivity and dimensionality needed to illuminate the spatiotemporal regulation of signaling networks in cells and tissues.

## Introduction

Protein kinases orchestrate diverse cellular processes by transmitting signals through site-specific phosphorylation of protein substrates. Genetically encoded fluorescent biosensors have illuminated these complex signaling networks by enabling direct and real-time monitoring of spatiotemporal kinase activities in living cells^1^. In particular, Förster resonance energy transfer (FRET)-based biosensors are the most well-developed and widely applied toolkit, providing a quantitative, ratiometric readout that reduces artifacts and signal fluctuations from variations in biosensor expression, photobleaching, or uneven illumination. Combined with subcellular targeting strategies, FRET-based biosensors have enabled the investigation of compartmentalized kinase signaling with high spatiotemporal resolution^2–6^.

Despite these advantages, most existing FRET-based biosensors use two spectrally distinct fluorescent proteins (FPs) as a FRET-pair, exhibiting limited dynamic range and sensitivity, which poses a challenge for detecting subtle changes in signaling activities^1^. Their large spectral footprints also make challenging to image multiple signaling activities via multiplexed activity imaging^7–9^. As the most commonly used donor, cyan fluorescent protein (CFP) also requires excitation with short-wavelength light, largely restricting the applicability in tissue or in vivo models^10^, and additionally exhibits complex, multiexponential fluorescence lifetime decays that complicate quantitative lifetime readout^11^. Although previous efforts have given rise to biosensors using FPs in longer wavelength windows, available far-red FP candidates often have suboptimal brightness and quantum yields, leading to compromised biosensor performance^12–15^. Chemigenetic biosensor designs, which integrate genetically encoded self-labeling protein tags with cell-permeant synthetic fluorophores, offer a promising strategy to overcome these limitations. Synthetic fluorophores, such as far-red rhodamines^16^ exhibit superior brightness, photostability, and spectral tunability, and can serve as genetically targetable fluorophores when used to label HaloTag^17^. This approach has been successfully applied to FRET-based biosensor development^18–23^. However, those chemigenetic designs have not been broadly applied to signaling biosensors, particularly kinase reporters, to enable highly sensitive detection of kinase activities.

Here, we report the development of a suite of highly sensitive chemigenetic FRET-based kinase biosensors with red-shifted emission, superior dynamic range and spectral versatility. These robust tools enable multiplexed activity imaging of GPCR signaling events and sensitive lifetime-based detection of kinase activity in acute somatosensory cortical slices, providing a versatile platform for dissecting the spatiotemporal regulation of complex signaling networks.

## Results

### Development and characterization of chemigenetic FRET-based AKAR

Previous studies showed that green fluorescent protein (GFP) forms a favorable interface with rhodamine-labeled HaloTag, achieving near-quantitative FRET efficiency^18^ ideal for developing biosensors with large dynamic ranges. Thus, we set out to overcome the limited dynamic range associated with current FRET-based kinase biosensors^1^ by incorporating a chemigenetic design^18–20^. Building upon the modular architecture of FRET-based kinase sensors, in which a phosphorylation-dependent molecular switch is sandwiched between a FRET-donor and FRET-acceptor^1^, we developed a chemigenetic sensor for cyclic adenosine monophosphate (cAMP)-dependent protein kinase (PKA) as an initial proof-of-concept. Specifically, we used a well-characterized switch comprising a PKA-specific substrate peptide and forkhead-associated 1 (FHA1) phospho-amino-acid binding domain (PAABD)^24,25^, separated by a 116-amino acid (116-aa) EV linker^26^, to promote conformational flexibility in the sensing unit. This switch was inserted between a FP FRET-donor and HaloTag covalently labeled with the far-red synthetic fluorophore JF_669_^16^ as the FRET-acceptor (Fig. 1a).

**Fig. 1:**
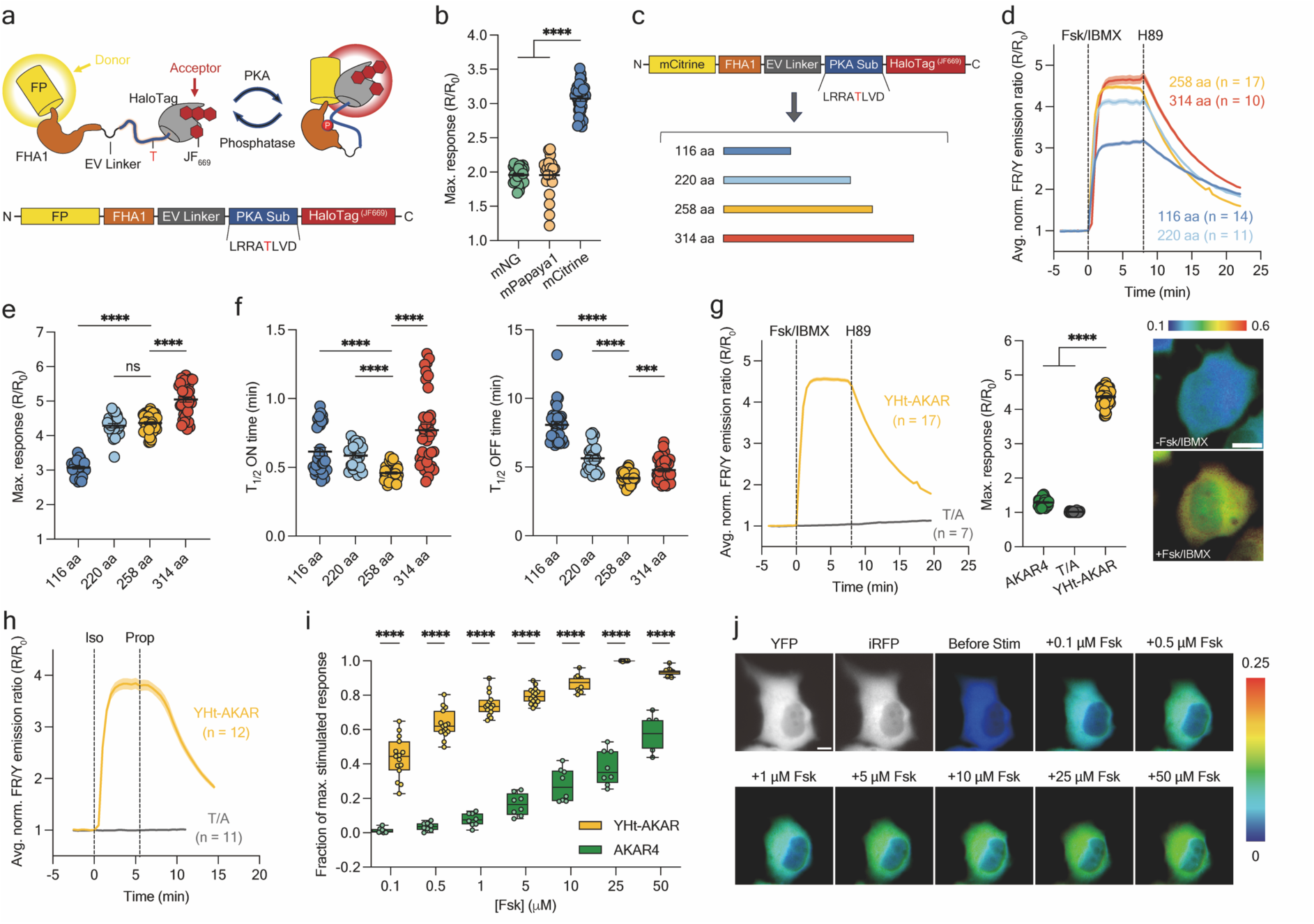
Design and characterization of YHt-AKAR. **a,** Schematic (top) and domain structure (bottom) of chemigenetic FRET-based AKAR containing FP donor, FHA1 domain, EV linker, PKA substrate, and HaloTag-JF_669_ acceptor. **b,** Maximum Fsk/IBMX-stimulated responses from sensor variants using mNG (n = 11), or mPapaya1 (n = 23), or mCitrine (n = 32) as the FRET-donor. **c,** Schematic of EV linker screening. **d,** Representative average time courses from HeLa cells expressing YHt-AKARs incorporating different EV linkers and treated with 50 μM Fsk/100 μM IBMX, followed by 20 μM H89. **e,** Maximum Fsk/IBMX-stimulated responses from YHt-AKAR EV linker candidates. n = 32 (116 aa), n = 23 (220 aa), n = 43 (258 aa), and n = 38 cells (314 aa) cells. **f,** Quantification of time to half-maximal response (T_1/2_ ON) after Fsk/IBMX addition (left) or time to half-maximal decay (T_1/2_ OFF) after H89 addition (right). **g,** Left, representative average time courses from HeLa cells expressing YHt-AKAR or YHt-AKAR (T/A) and treated with Fsk/IBMX then H89. Middle, maximum Fsk/IBMX-stimulated responses from AKAR4 (n = 31), YHt-AKAR (T/A) (n = 26), and YHt-AKAR (n = 43). Right, representative pseudocolor images of HeLa cells expressing YHt-AKAR labeled with JF_669_ before and after Fsk/IBMX stimulation. Warmer colors indicate higher ratios. Scale bar, 10 μm. **h,** Representative average time courses of HEK293T cells expressing YHt-AKAR and treated with 1 μM isoproterenol (Iso) and 10 μM propranolol (Prop). **i,** Concentration-response measurements in HeLa cells expressing either YHt-AKAR or AKAR4. Cells were stimulated with the indicated concentrations of Fsk, followed by stimulation with 100 μM IBMX. Data are plotted as (ΔR[Fsk])/(ΔR_max_), box-and-whisker plots showing the median, interquartile range, minimum, maximum and mean. Statistical analyses were performed using unpaired, two-tailed Welch’s unequal variance t-test, \*\*\*\**P* < 0.0001. n = 8 (AKAR4), and n = 15 (YHt-AKAR). **j,** Representative YFP and iRFP widefield images, and FR/Y pseudocolor images showing the responses of YHt-AKAR to different Fsk concentration stimulation in HeLa cells. Time courses are representatives of three independent experiments. Dashed lines indicate addition of drug, solid lines indicate mean responses, and shaded areas depict SEM. For maximum response measurements, individual data points are pooled from three independent experiments. Error bars depict mean ± SEM. Data in b and e-g were analyzed using one-way ANOVA followed by Dunnett’s multiple-comparison test, \*\*\*\**P* < 0.0001, \*\*\**P* = 0.0004, ns = not significant.

We first tested three candidates to identify an optimal donor FP to pair with HaloTag-JF_669_: mNeonGreen (mNG)^27^, a bright FP with similar spectral characteristics as GFP, as well as mPapaya1^28^ and mCitrine^29^, two bright yellow-emitting FPs. HeLa cells transiently expressing each candidate sensor and loaded with JF_669_ were stimulated with the adenylyl cyclase activator forskolin (Fsk) and phosphodiesterase inhibitor 3-isobutyl-1-methylxanthine (IBMX) to elicit maximal PKA activation and sensor response, which we monitored as the change in the acceptor-to-donor emission ratio (Δ*R/R_0_*). The mCitrine-containing sensor showed the largest response to maximal PKA stimulation (i.e., dynamic range) (Δ*R/R*_0_ = 207.3% ± 3.9%, mean ± SEM, n = 32 cells) (Fig. 1b) and was therefore selected for further development.

We next set out to optimize the linker separating the PKA substrate and FHA1 domain. Incorporating an EV linker increases separation between the donor and acceptor fluorophores in the sensor off state, thereby minimizing basal FRET and enhancing biosensor dynamic range^26^, yet previous studies have differed in their selection of the optimal EV linker length^26,30^. Thus, we compared the effects of four EV linker lengths: 116 aa, 220 aa, 258 aa, and 314 aa (Fig. 1c). As expected, we observed progressively lower basal far-red over yellow (FR/Y) emission ratios (R_0_) with increasing EV linker length, reflecting reduced basal proximity between the donor and acceptor fluorophores (Supplementary Fig. 1). Overall, we also observed progressively higher stimulated ratios as EV linker length was increased, with the 314 aa linker ultimately yielding the largest dynamic range (Δ*R/R*_0_ = 404.7% ± 7.6%, n = 38 cells) (Fig. 1d-e). However, comparing the response kinetics revealed that the 258-aa variant has the fastest time to half-maximal response (t_1/2_ ON = 0.46 ± 0.01 min, n = 43 cells), as well as the fastest half-maximal decay time upon treatment with the PKA inhibitor H89 (20 μM) to reverse the sensor response (t_1/2_ OFF = 4.19 ± 0.06 min, n = 43 cells). Alternatively, the 314-aa variant yielded significantly slower kinetics (t_1/2_ ON = 0.77 ± 0.04 min, *P* < 0.0001; t_1/2_ OFF = 4.78 ± 0.13 min, *P* < 0.001, n = 38 cells) (Fig. 1f). Considering the balance between dynamic range and response kinetics, we therefore selected the 258-aa linker variant as the optimal configuration, which we designated as yellow/far-red HaloTag A kinase activity reporter (YHt-AKAR).

YHt-AKAR displayed an emission ratio change of Δ*R/R*_0_ = 336.6% ± 3.7% (n = 43 cells) upon Fsk/IBMX stimulation in HeLa cells, followed by rapid reversal upon H89 addition (Fig. 1g). Meanwhile, a Thr-to-Ala (T/A) phosphor-acceptor mutant sensor showed no detectable response to Fsk/IBMX stimulation, confirming that YHt-AKAR emission ratio changes were due to PKA-mediated phosphorylation (Fig. 1g). In HEK293T cells, stimulation of endogenous β_2_-adrenergic receptors using 1 μM isoproterenol similarly elicited a rapid rise in YHt-AKAR emission ratio (Δ*R/R*_0_ = 237.3% ± 13.2%, n = 22 cells) that was promptly reversed by addition of the β-blocker propranolol (10 μM), while the T/A mutant remained unresponsive (Fig. 1h). Remarkably, the Fsk/IBMX-induced YHt-AKAR exhibits an approximately 7-fold higher dynamic range compared with AKAR4^7^ (Fig. 1g), and concentration-response assays revealed that YHt-AKAR responds to Fsk concentrations as low as 0.1 μΜ, whereas AKAR4 only responded at 10-fold higher concentrations (i.e., 1 μM) (Fig. 1i-j). YHt-AKAR also reached saturation near 25 μM Fsk, while AKAR4 remained submaximal at 50 μM, indicating substantially higher sensitivity of YHt-AKAR. These results establish YHt-AKAR as a robust and highly sensitive chemigenetic tool for monitoring PKA activity in living cells.

### Extending the chemigenetic FRET design to red-shifted donors

Red-shifted sensors offer key advantages for advanced imaging applications, including flexibility in multiplexed imaging, reduced interference by cellular autofluorescence, and enhanced tissue penetration. We next sought to extend the chemigenetic FRET design to incorporate a red-shifted donor. Despite the robust performance of YHt-AKAR, it is unclear whether RFPs, with their distinct species origin, would demonstrate comparable performance as a GFP-derived FRET-donor when paired with HaloTag-JF_669_. To investigate this possibility, we screened a library of ten candidate RFPs (sTagRFP^31^, mRuby3^32^, Azalea B5^33^, mNeptune2^34^, mKoκ^35^, FusionMQV^36^, FusionRed^37^, mScarlet-I^38^, mScarlet^39^, and mCherry^40^) in place of mCitrine in the YHt-AKAR backbone to identify a suitable FRET-donor (Fig. 2a). As above, we expressed each biosensor variant in HeLa cells labeled with JF_669_ and stimulated with Fsk/IBMX to achieve maximal PKA activation. Under these conditions, the sTagRFP-containing sensor exhibited the largest far-red over red (FR/R) emission ratio change (Δ*R/R*_0_ = 114.2% ± 2.0%, n = 25 cells), significantly outperforming all other RFP variants (Fig. 2b-c) and establishing sTagRFP as the optimal FRET-donor for subsequent development of chemigenetic RHt-sensors (R/FR).

**Fig. 2:**
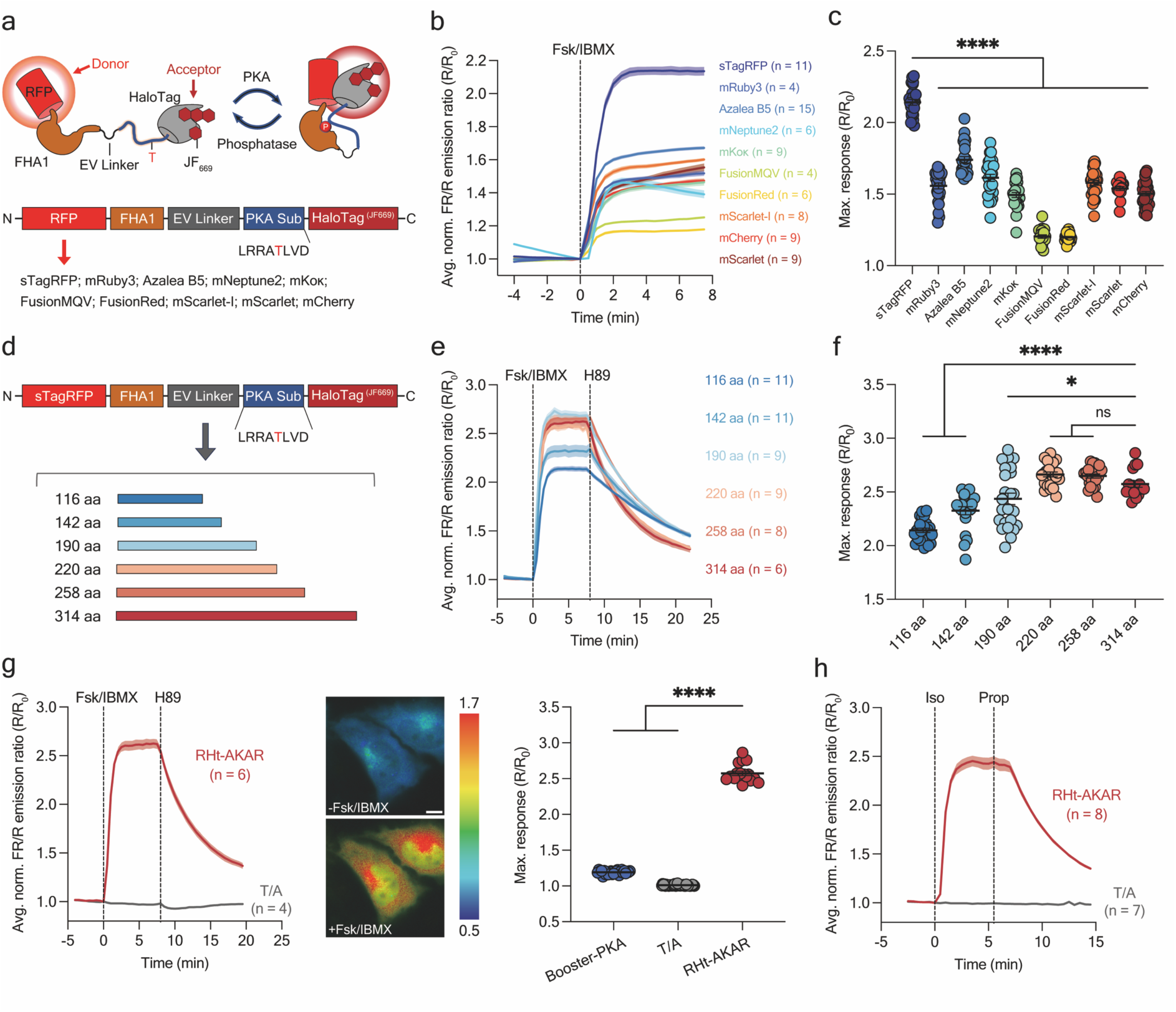
Design and characterization of RHt-AKAR. **a,** Schematic (top) and domain structure (bottom) of RHt-AKAR. **b,** Representative average time courses of HeLa cells expressing RHt-AKAR variants incorporating the indicated RFP donors and treated with 50 μM Fsk/100 μM IBMX. **c,** Maximum Fsk/IBMX-stimulated responses for RHt-AKAR donor variants. n = 25 (sTagRFP), n = 28 (mRuby3), n = 25 (Azalea B5), n = 27 (mNeptune2), n = 26 (mKoκ), n = 20 (FusionMQV), n = 22 (FusionRed), n = 29 (mScarlet-I), n = 17 (mScarlet), and n = 35 cells (mCherry). **d,** Scheme illustrating the EV linker screen. **e,** Representative average time courses of HeLa cells expressing RHt-AKAR EV linker candidates and treated with Fsk/IBMX and 20 μM H89. **f,** Maximum Fsk/IBMX-stimulated responses from RHt-AKAR EV linker variants. n = 25 (116 aa), n = 22 (142 aa), n = 25 (190 aa), n = 24 (220 aa), n = 23 (258 aa), and n = 17 cells (314 aa). **g,** Left, representative average time courses of HeLa cells expressing RHt-AKAR or RHt-AKAR (T/A) and treated with Fsk/IBMX and subsequently H89. Middle, representative FR/R pseudocolor images of HeLa cells expressing RHt-AKAR labeled with JF_669_ before and after Fsk/IBMX stimulation. Warmer colors indicate higher ratios. Scale bar, 10 μm. Right, maximum Fsk/IBMX-stimulated responses from HeLa cells expressing Booster-PKA (n = 12), RHt-AKAR (T/A) (n = 38), or RHt-AKAR (n = 17 cells). **h,** Representative average time courses of HEK293T cells expressing RHt-AKAR and treated with 1 μM isoproterenol (Iso) then 10 μM propranolol (Prop). Time courses are representatives of three independent experiments. Dashed lines indicate addition of drug, solid lines indicate mean responses, and shaded areas depict SEM. For maximum response measurements, individual data points are pooled from three independent experiments. Error bars indicate mean ± SEM. Statistical analyses were performed using one-way ANOVA followed by Dunnett’s multiple-comparison test, \*\*\*\**P* < 0.0001, \**P* = 0.0327, ns = not significant.

To further enhance the dynamic range, we expanded the EV linker library to 6 lengths (116 aa, 142 aa, 190 aa, 220 aa, 258 aa, and 314 aa) to verify the optimal linker length for R/FR FRET-pair (Fig. 2d), given potential interface differences between RFPs-donor and HaloTag. In JF_669_-loaded HeLa cells expressing the corresponding biosensors, we again observed a clear decrease in the basal emission ratio (R_0_) with increasing EV linker length, similar to our YHt-AKAR results (Supplementary Fig. 2). In contrast to YHt-AKAR, however, stimulation with Fsk/IBMX followed by H89 revealed that the 314-aa linker variant exhibited the best combination of dynamic range and response kinetics (Fig. 2e-f and Supplementary Fig. 3), highlighting the importance of tailoring linker length for individual FRET-pairs. Specifically, the variant, named RHt-AKAR, showed a 157.4% ± 3.0% (n = 17 cells) increase in far-red over red (FR/R) emission ratio upon maximal stimulation of PKA with Fsk/IBMX, whereas RHt-AKAR (T/A) showed no detectable response to Fsk/IBMX stimulation (Fig. 2g). RHt-AKAR also showed robust performance in Iso-stimulated HEK293T cells (Δ*R/R*_0_ = 120.6% ± 8.8%, n = 16 cells) (Fig. 2h). Compared with the current best-in-class red-shifted PKA sensor Booster-PKA^30^, RHt-AKAR exhibits approximately a 7-fold higher dynamic range (Fig. 2g). Thus, RHt-AKAR is a highly sensitive chemigenetic PKA sensor and represents a dramatic improvement over existing sensors in the red/far-red spectral range. Our results also demonstrate that the chemigenetic FRET platform is broadly adaptable across diverse spectral regions and FP families.

### Generalizing the chemigenetic FRET design to other signaling activities

Many FP-based FRET sensors are limited by their poor dynamic range and narrow spectral window. The design of our chemigenetic Ht-AKARs retains the characteristic modular architecture of FRET-based sensors, in which the sensing unit can be exchanged to detect various signaling events^1^, providing a robust foundation to expand this design beyond PKA to build a versatile platform for probing diverse signaling events underlying various cellular functions.

We first sought to generalize the chemigenetic FRET design to another AGC-family kinase. Specifically, we adapted the Ht-AKARs framework to protein kinase C (PKC)^41^ by replacing the PKA substrate sequence with a PKC substrate^41^ to obtain both Y/FR and R/FR HaloTag C kinase activity reporters (i.e., YHt-CKAR and RHt-CKAR) (Fig. 3a-b). In HeLa cells, stimulation with 50 ng/mL phorbol 12-myristate 13-acetate (PMA) triggered prominent, 140.9% ± 3.5% (n = 44 cells) and 72.0% ± 1.7% (n = 19 cells) emission ratio increases from YHt-CKAR and RHt-CKAR, respectively (Fig. 3a-b), representing the largest dynamic ranges achieved thus far using FRET-based PKC reporters^42,43^. In contrast, T/A phosphorylation-site mutants showed no response to PMA stimulation (Fig. 3a-b), verifying sensor specificity for PKC activity.

**Fig. 3:**
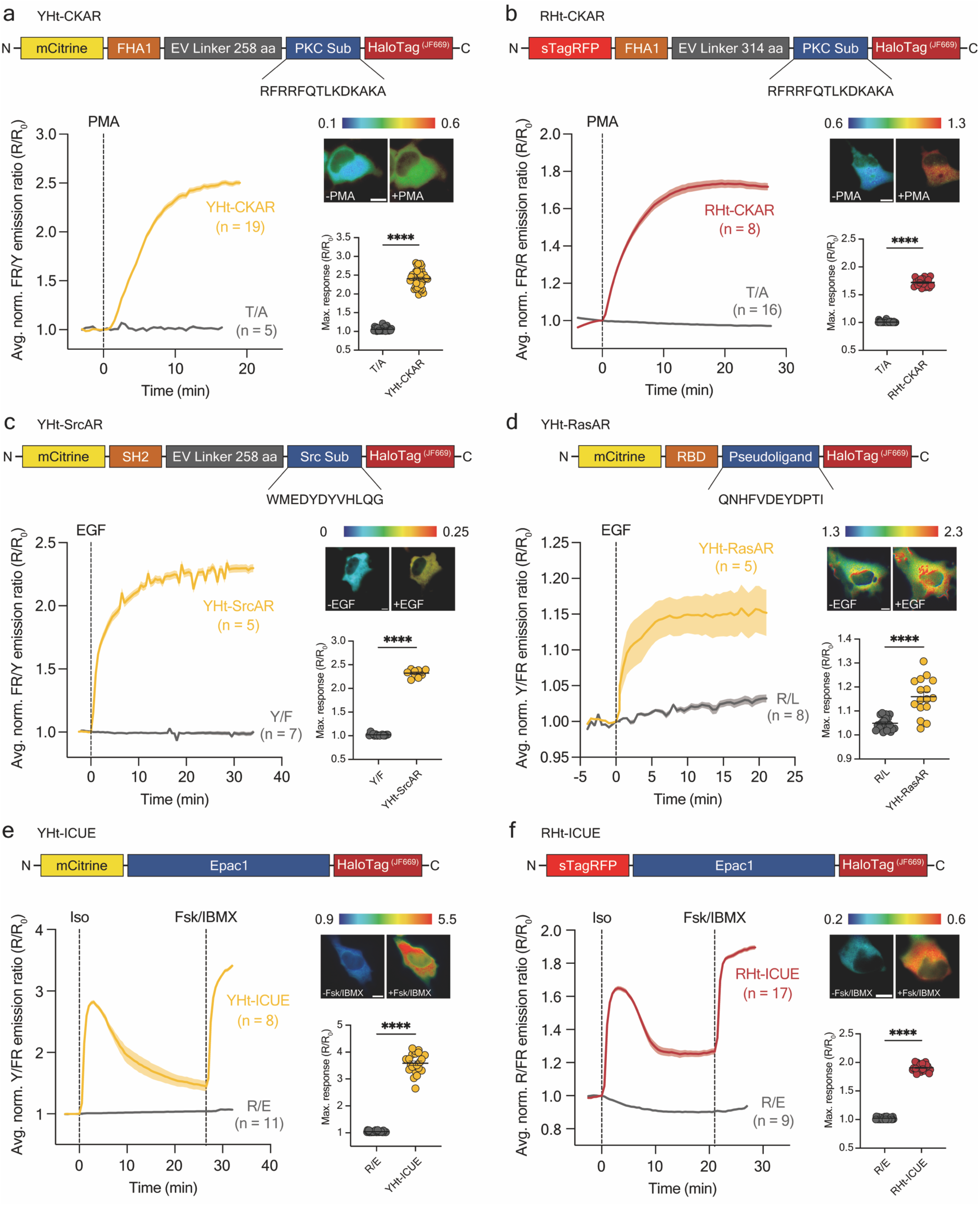
Construction of chemigenetic biosensors for PKC, Src, Ras, and cAMP based on Ht-AKARs design. **a-f,** Top, domain structures of YHt-ICUE **(a)**, RHt-ICUE **(b)**, YHt-SrcAR **(c)**, YHt-RasAR **(d)**, YHt-ICUE **(e)**, and RHt-ICUE **(f)**. Bottom left, representative average time courses of HeLa cells expressing **(a)** YHt-CKAR or YHt-CKAR (T/A), or RHt-CKAR or RHt-CKAR (T/A) **(b)** and treated with 50 ng/mL PMA; COS-7 cells expressing **(c)** YHt-SrcAR or YHt-SrcAR (Y/F) or **(d)** YHt-RasAR or YHt-RasAR (R/L), and treated with 100 ng/mL EGF; **(e)** HEK293T cells expressing YHt-ICUE or YHt-ICUE (R/E) or **(f)** RHt-ICUE or RHt-ICUE (R/E) and treated with 1 μM isoproterenol (iso) and 50 μM Fsk/100 μM IBMX. Upper right, representative pseudocolor images of HeLa cells expressing **(a)** YHt-CKAR or **(b)** RHt-CKAR before and after PMA stimulation, **(c)** COS-7 cells expressing YHt-SrcAR, or **(d)** YHt-RasAR before and after EGF stimulation, or HEK293T cells expressing **(e)** YHt-ICUE, or **(f)** RHt-ICUE before and after Fsk/IBMX stimulation. Warmer colors indicate higher ratios. Scale bars, 10 μm. Bottom right, maximum PMA-stimulated responses from **(a)** YHt-CKAR (n = 44) and YHt-CKAR (T/A) (n = 26), or **(b)** RHt-CKAR (n = 19) and RHt-CKAR (T/A) (n = 40), EGF-stimulated responses from **(c)** YHt-SrcAR (n = 10) and YHt-SrcAR (Y/F) (n = 15) or **(d)** YHt-RasAR (n = 16), YHt-RasAR (R/L) (n = 23), or Fsk/IBMX-stimulated responses from **(e)** YHt-ICUE (n = 22) and YHt-ICUE (R/E) (n = 33), or **(f)** RHt-ICUE (n = 31), RHt-ICUE (R/E) (n = 30). Time courses are representatives of three independent experiments. Dashed lines indicate addition of drug, solid lines indicate mean responses, and shaded areas depict SEM. For maximum response measurements, individual data points are pooled from three independent experiments. Error bars depict mean ± SEM. Statistical analyses were performed using unpaired, two-tailed Student’s t-tests, \*\*\*\**P* < 0.0001.

Next, we applied our chemigenetic platform to the proto-oncogenic protein tyrosine kinase Src (Src)^44^, which requires not only incorporating a new substrate sequence but also swapping the FHA1 PAABD for a Src homology 2 (SH2) domain^45^ (Fig. 3c). The conformational change induced by this sensing domain typically yields much smaller responses from current FRET-based Src activity reporters (SrcARs)^31,44–47^ compared with similar FHA1-containing sensors. However, COS-7 cells expressing YHt-SrcAR exhibited a remarkable 132.3% ± 2.4% (n = 10 cells) emission ratio increase upon stimulation with 100 ng/mL EGF (Fig. 3c), offering a substantial improvement over existing biosensors for monitoring Src activity and establishing the application of this chemigenetic platform beyond serine/threonine kinases.

Besides kinases, we were also able to successfully adapt our chemigenetic approach to sense the activity of the small GTPase Ras^48^. Our design was based on a recently developed FRET-based Ras activity reporter (RasAR), which features a bipartite sensing unit comprising the Raf1 Ras-binding domain (RBD) and an RBD-interacting pseudoligand derived from Ras^48^. We thus constructed a Y/FR-HaloTag-Ras activity reporter (YHt-RasAR) by replacing the C/Y FP pair in RasAR with mCitrine/HaloTag-JF_669_ (Fig. 3d). In COS-7 cells expressing a dominant negative HRas(S17N), which provides a low-activity baseline by disrupting Ras-RBD engagement, YHt-RasAR showed an average Y/FR emission ratio of 1.70 ± 0.03 (n = 149 cells), while cells expressing constitutively active HRas (G12V) yielded a ratio of 2.55 ± 0.05 (n = 104 cells), translating into a 50.00% ± 0.05% dynamic range (R[CA]-R[DN])/R[DN]; Supplementary Fig. 4). Furthermore, YHt-RasAR reported a 16.0% ± 1.9% (n = 16 cells) increase in the Y/FR emission ratio upon EGF stimulation of the Ras signaling pathway in COS-7 cells overexpressing WT HRas (Fig. 3d).

Finally, we applied our chemigenetic FRET platform to monitor the second messenger cAMP by developing HaloTag-based indicators of cAMP using Epac1 (Ht-ICUEs) (Fig. 3e-f). Specifically, we inserted the Epac1-based sensing domain from the C/Y-FRET sensor ICUE4^9^ between our Y/FR and R/FR chemigenetic FRET-pairs (Fig. 3e-f). In HEK293T cells, YHt-ICUE and RHt-ICUE showed large, transient responses of 182.4% ± 3.2% (n = 22 cells) and 65.1% ± 1.3% (n = 31 cells), respectively, upon Iso stimulation followed by larger, sustained 258.6% ± 8.1% (n = 22 cells) and 90.5% ± 1.0% (n = 31 cells) ratio increases, respectively, upon subsequent Fsk/IBMX treatment (Fig. 3e-f). Conversely, cAMP-binding-deficient R279E (R/E) mutant sensors failed to respond to Iso or Fsk/IBMX stimulation. Together, these results demonstrate the generalizability of our chemigenetic FRET platform, which can report diverse classes of signaling events with high dynamic range and specificity.

### Multiplexed activity imaging reveals unexpected GPCR isoform selective signaling

Multiplexed biosensor readouts enable simultaneous, real-time dissection of signaling network interactions in single living cells. Our chemigenetic FRET-based sensors, with their large dynamic ranges and distinct spectral range, should serve as robust tools for more sensitive multiplexed detection of signaling molecules alongside existing reporters. Indeed, when we co-expressed RHt-AKAR with the green, excitation-ratiometric PKC sensor ExRai-CKAR^49^ in HEK293T cells to track PKA and PKC dynamics, we observed a large PMA-induced ExRai-CKAR response (Δ*R/R*_0_ = 59.3% ± 4.4%, n = 12 cells) that was rapidly reversed by addition of the pan-PKC inhibitor Gö6983 (10 μM) (Fig. 4a), while RHt-AKAR specifically responded with a large ratio increase (Δ*R/R*_0_ = 107.5% ± 8.3%, n = 12 cells) upon Fsk/IBMX addition, which was reversed by H89 treatment (Fig. 4a). FRET-based sensors with the same acceptor but spectrally distinct donors can also be readily multiplexed ^50^, and we were able to perform concurrent tracking of intracellular cAMP levels and PKA activity in HEK293T cells co-expressing YHt-ICUE and RHt-AKAR. Both sensors showed large Fsk/IBMX-induced emission ratio changes (Δ*R/R*_0_ = 145.1% ± 2.8% for RHt-AKAR and 105.8% ± 5.9% for YHt-ICUE, n = 20 cells), with H89 treatment selectively reversing the RHt-AKAR response without affecting the YHt-ICUE signal (Fig. 4b). These results establish our chemigenetic FRET sensors as a robust platform for sensitive multiplexed activity imaging alongside both FP-based and chemigenetic sensors.

**Fig. 4:**
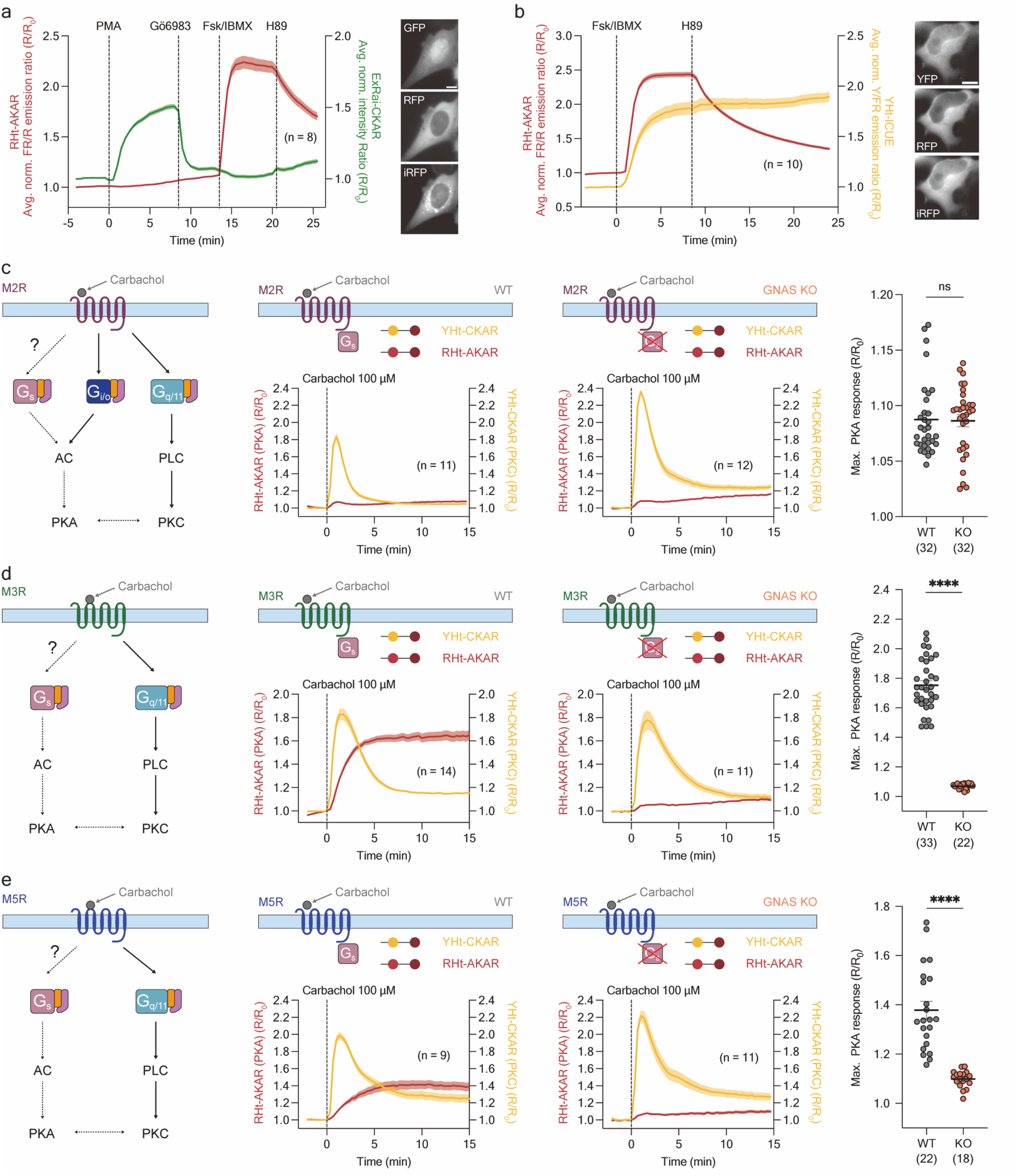
Multiplexed activity imaging using chemigenetic biosensors. **a-b,** Left, representative average time courses of HeLa cells expressing RHt-AKAR (red) and ExRai-CKAR (green) and treated with 50 ng/mL PMA, 10 μM Gö6983, 50 μM Fsk/100 μM IBMX and 20 μM H89 **(a)**, or HEK293T cells expressing YHt-ICUE (yellow) and RHt-AKAR (red) and treated with Fsk/IBMX and H89 **(b)**. Right, representative epifluorescence images of each channel. Scale bars, 10 μm. **c-e,** Far left, schematic illustration of signaling downstream of M2R **(c)**, or M3R **(d)**, or M5R **(e)**. Left, representative average time courses of HEK293A cells (parental) expressing RHt-AKAR (red), YHt-CKAR (yellow), plus M2R **(c)**, or M3R **(d)**, or M5R **(e)**, and treated with 100 μM carbachol. Right, average time courses of HEK293A GNAS KO cells expressing RHt-AKAR (red), YHt-CKAR (yellow), and M2R **(c)**, or M3R **(d)**, or M5R **(e)** and treated with 100 μM carbachol. Far right, maximum PKA responses induced within 5 min of 100 μM carbachol addition mediated by M2R in HEK293A (parental, n = 32) or HEK293A GNAS KO (n = 32) cells **(c)**, M3R in HEK293A (parental, n = 33) or HEK293A GNAS KO (n = 22) cells **(d)**, or M5R in HEK293A (parental, n = 22) or HEK293A GNAS KO (n = 18) cells **(e)**. Time courses are representatives of three independent experiments. Dashed lines indicate addition of drug, solid lines indicate mean responses, and shaded areas depict SEM. For maximum response measurements, individual data points are pooled from three independent experiments. Error bars show mean ± SEM. Statistical analyses were performed using unpaired, two-tailed Student’s t-tests, \*\*\*\**P* < 0.0001.

Next, we investigated the signaling architecture mediated by distinct G protein classes downstream of different GPCR isoforms. Muscarinic acetylcholine receptors (mAChRs) are major neuromodulatory GPCRs that transduce acetylcholine signals to regulate numerous physiological processes, including heart rate, smooth muscle contraction, and neuronal excitability^51^. Prior work showed that M_1_ muscarinic receptor (M1R) can engage PKA through Ca^2+^- or PKC-dependent crosstalk downstream of G_q/1152_, but whether other mAChR isoforms display comparable PKA engagement has remained unclear. To investigate isoform-specific coupling in a unified assay, we performed multiplexed activity imaging using RHt-AKAR to report PKA activity as a G_s_ readout and YHt-CKAR to monitor G_q_-induced PKC activity upon carbachol stimulation. Within this framework, we examined three isoforms with distinct canonical couplings: M_2_ muscarinic receptor (M2R), a classically G_i/o_- and G_q/11_-coupled receptor known to inhibit adenylyl cyclase while activating PKC^53,54^ (Fig. 4c), and both M_3_ (M3R) and M_5_ muscarinic receptors (M5R), which are canonically coupled to only G_q/11_^54,55^ (Fig. 4d-e).

Upon 100 μM carbachol stimulation, M2R-expressing HEK293A cells showed an immediate, transient YHt-CKAR response (Δ*R/R*_0_ = 67.2% ± 4.1%, n = 32 cells) with only a small, gradual ratio increase (Δ*R/R*_0_ = 8.7% ± 0.6%, n = 32 cells) from RHt-AKAR (Fig. 4c). Small PKA responses were observed only at 100 μM or higher carbachol concentrations (Supplementary Fig. 5), suggesting weak activation of PKA signaling. By contrast, M3R-expressing cells exhibited clear and sustained RHt-AKAR responses (Δ*R/R*_0_ = 75.3% ± 3.1% n = 33 cells), together with transient YHt-CKAR responses (Δ*R/R*_0_ = 67.7% ± 3.6%, n = 33 cells) under the same stimulation conditions (Fig. 4d), with both sensors showing robust responses across a range of carbachol concentrations (Supplementary Fig. 6). M3R-mediated PKA activation was further confirmed under endogenous receptor expression conditions (predominantly M3R) in HEK293A cells transfected with the ultrasensitive reporter HaloAKAR^56^ (Supplementary Fig. 7). HEK293A cells overexpressing M5R produced similar downstream responses to those observed with M3R (PKA Δ*R/R*_0_ = 41.2% ± 3.6%; PKC Δ*R/R*_0_ = 90.4% ± 3.5%, n = 22 cells) (Fig. 4e). Concentration-response assays further revealed that the M5R-mediated PKA response can be detected at carbachol concentrations higher than 1 μM (Supplementary Fig. 8). These experiments demonstrate PKC activation across mAChR isoforms and reveal robust PKA engagement with M3R and M5R but not M2R.

Consistent with their reported G_q/11_ coupling, pretreatment with the G_q/11_ blocker FR900359 suppressed the carbachol-induced YHt-CKAR responses from all three receptors (Supplementary Fig 9). Notably, the weak M2R-mediated PKA response was also completely suppressed by FR900359 pretreatment (Supplementary Fig. 9), but was unaffected by knocking out GNAS, which encodes the G_s_ subunit (Fig. 4c), or by G_i/o_ blockade via pretreatment with pertussis toxin (PTX) (Supplementary Fig. 9), suggesting that cAMP/PKA pathway cross-activation is driven by G_q/11_ signaling^52^. However, FR900359 or PTX pretreatment had no effect on the carbachol-stimulated RHt-AKAR responses observed in M3R- or M5R-overexpressing HEK293A cells (Supplementary Fig. 9), which were instead nearly abolished in GNAS knockout cells (M3R Δ*R/R*_0_ = 9.1% ± 0.6%, n = 22 cells; M5R Δ*R/R*_0_ = 11.8% ± 0.8%, n = 18 cells) (Fig. 4d-e). Multiplexed activity imaging of YHt-ICUE and RHt-AKAR in M3R-overexpressing cells further showed rapid and sustained increases in both cAMP and PKA upon carbachol (100 μM) that were abolished in the GNAS knockout context, confirming G_s_-adenylyl cyclase-mediated cAMP/PKA activation (Supplementary Fig. 10). Together, these results suggest that M3R and M5R couple to G_s_ to activate PKA signaling, in addition to their classical G_q/11_-PKC signaling. Our findings support a previous study showing that M3R can signal through multiple G protein families including G_s_ and demonstrate M5R exhibits similar promiscuous coupling^57^. Our study further highlights isoform-selective signaling architecture, in which individual receptor subtypes recruit distinct G protein combinations to shape unique downstream signaling networks.

### RHt-AKAR enables robust PKA activity imaging in brain slices

PKA is a central effector of neuromodulator pathways in the brain, where dopamine, norepinephrine, and acetylcholine regulate synaptic plasticity, excitability, and network states via GPCR-mediated signaling^58,59^. PKA phosphorylates key substrates within axons and dendritic spines, where A kinase anchoring protein (AKAP)-mediated scaffolding creates PKA activity microdomains^60^. The ability to monitor subtle and rapid activity changes associated with these domains is essential for revealing how complex neurological processes are regulated in the brain. It requires sensors with high dynamic range and optimal spectral properties for imaging in complex tissues. Two-photon fluorescence lifetime imaging microscopy (2p-FLIM) is widely employed to perform quantitative imaging of FRET-based sensors with high spatial resolution in scattering tissues^61,62^. The red-emitting donor and high dynamic range of RHt-AKAR should be ideally suited for monitoring PKA activity via 2p-FLIM FRET imaging. Indeed, HEK293T cells expressing RHt-AKAR exhibited a significant decrease in donor lifetime (− Δlifetime = 0.185 ns ± 0.046 ns, n = 6 cells) upon Fsk stimulation (Fig. 5a-b). Notably, the far-red fluorophore JF669 is cell-permeant and bioavailable *in vivo*, supporting the applicability of these chemigenetic FRET sensors to deep-tissue and physiological imaging^16^. Thus, we next tested whether RHt-AKAR can reveal endogenous GPCR-mediated signaling in brain tissue. In acute mouse somatosensory cortical slices pre-incubated with JF_669_, RHt-AKAR expressed well in neurons and clearly reported β-adrenergic activation with highly sensitive lifetime decrease (Fig. 5c). Stimulation with 1 μM Iso induced a reproducible drop in lifetime relative to baseline (−Δlifetime = 0.224 ns ± 0.011 ns, n = 7 neurons from 3 animals), which further decreased with subsequent Fsk treatment (−Δlifetime = 0.356 ns ± 0.014 ns, n = 7 neurons from 3 animals) (Fig. 5c-d). RHt-AKAR thus exhibits a larger dynamic range than previously reported signaling lifetime biosensors, including the commonly used FLIM-AKAR^63^.

**Fig. 5:**
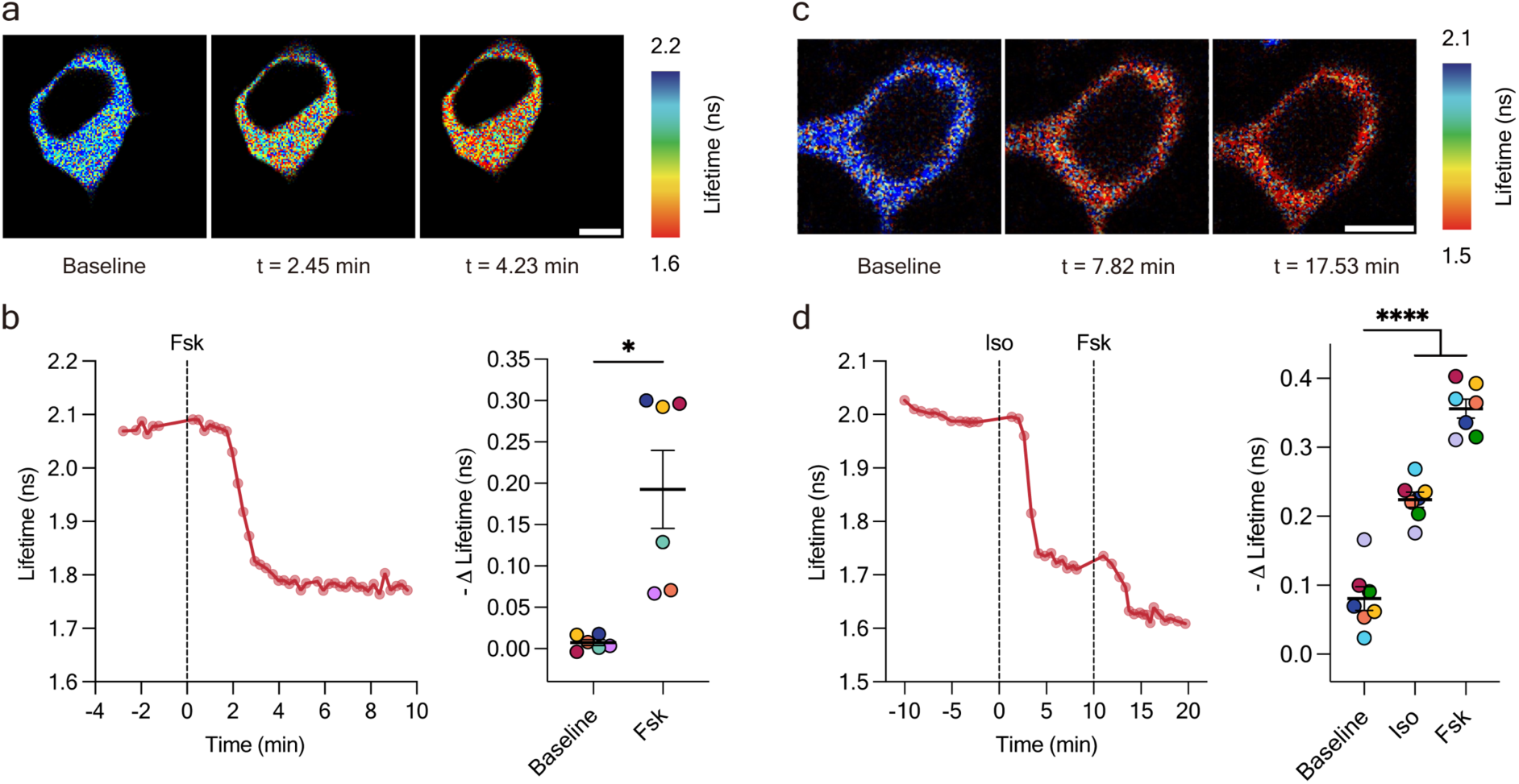
2p-FLIM imaging of RHt-AKAR in cultured cells and brain slices. **a,** Representative pseudocolor images of HEK293T cells expressing RHt-AKAR labeled with JF_669_ before (baseline) and after (t = 2.45 min, or t = 4.23 min) 50 μM Fsk stimulation. **b,** Left, representative single-cell lifetime trace from a HEK293T cell expressing RHt-AKAR and treated with 50 μM Fsk. Right, quantification of maximum lifetime change (n = 6 cells). **c,** Representative pseudocolor images of an excitatory neuron in an acute cortical slice expressing RHt-AKAR labeled with JF_669_ before (baseline) and after 1 μM isoproterenol (Iso) stimulation (t = 7.82 min) later supplemented with 50 μM Fsk (t = 17.53 min), **d,** Left, representative single-cell lifetime trace from an excitatory neuron in an acute cortical slice expressing RHt-AKAR and treated with 1 μM isoproterenol and 50 μM Fsk. Right, quantification of maximum lifetime change (n = 7 neurons from 3 animals). Data points with the same color are from the same cell. \**P* = 0.031, paired Wilcoxon signed rank test; \*\*\*\**P* < 0.0001, non-parametric Kruskal-Wallis’s test followed by Dunnett’s multiple-comparison test. Scale bars, 10 μm.

These results establish RHt-AKAR as a powerful tool for deep-tissue kinase activity imaging, providing a sensitive, multiplexing-compatible platform for mapping subtle, physiologically relevant signaling dynamics *in vivo*. Furthermore, our work demonstrates the versatility and utility of this chemigenetic FRET-based biosensor platform, which offers robust FRET responses with spectral tunability and enables sensitive visualization of different signaling activities with superior performance.

## Discussion

Precise modulation of kinase signaling is fundamental for regulating diverse cellular functions, from hormone secretion^64,65^ to gene expression^66–68^ and stress responses^69,70^. However, such subtle changes often approach the detection limits of conventional, entirely FP FRET-based biosensors, making them difficult to capture with sufficient sensitivity and precision. Here, our work demonstrates that novel chemigenetic biosensors, which incorporate HaloTag labeled with the far-red synthetic fluorophore JF_669_, markedly expand the capability to visualize these biologically important signaling events via quantitative FRET-based readouts. In particular, Ht-AKARs exhibited substantially improved dynamic ranges compared to traditional FP-based designs^7,30^ and enabled detection of rapid, small-amplitude PKA activity shifts that would otherwise remain obscured. These findings underscore the ability of our chemigenetic tools to illuminate fine-tuned aspects of kinase regulation in living cells. By exploiting the modularity of Ht-AKARs, we also successfully extended this chemigenetic FRET design to generate biosensors for monitoring PKC, Src, Ras, and cAMP dynamics, establishing the generalizability of this chemigenetic platform for developing high-performance, multiplexable biosensors to monitor various signaling activities.

Physiological processes are rarely driven by individual signaling pathways but instead emerge from the interplay of intricate signaling networks acting within spatially compartmentalized microdomains^49,56^. Using our chemigenetic biosensors, we were able to achieve robust multiplexed activity imaging to simultaneously monitor distinct signaling activities within single living cells, enabling direct access of pathway-specific signaling architectures downstream of distinct mAChR isoforms. M3R and M5R were shown to display non-canonical G_s_-mediated PKA activation in response to carbachol stimulation, whereas M2R exhibited weak, G_q/11_-dependent PKA crosstalk under the same condition. These results highlight the strength of this chemigenetic platform to quantify receptor-defined signaling states and isoform-specific patterns within a single receptor family. By preserving high sensitivity during multiplexed readouts, these tools open the door to uncovering subtle variations in signaling network behavior that were previously inaccessible^31,49^, thereby advancing opportunities to decode the dynamic signaling interplay and crosstalk that underlie cellular function. Furthermore, biological tissue imaging of kinase dynamics is essential for bridging cellular mechanisms with physiological and pathological processes, yet achieving sufficient sensitivity and spectral performance in native tissues has remained a major challenge^71^. Here, we established the utility of RHt-AKAR for tissue imaging, taking advantage of the red-shifted spectral properties that provide superior biological tissue penetration and reduced autofluorescence signal. Two-photon FLIM characterization revealed that RHt-AKAR achieves a markedly higher performance compared with the current state of the art^63^. Importantly, the compatibility of this tool with FLIM enables quantitative, intensity-independent measurements, allowing robust comparison of kinase activity across cells, animals, and experimental time scales. Notably, this robust tool can be applied to report endogenous GPCR activation in acute brain slices, underscoring its compatibility with complex and physiologically relevant signaling pathways. By monitoring kinase activity with high fidelity in brain tissue, these chemigenetic biosensors initiate new possibilities for dissecting how dynamic signaling perturbations reshape cellular functions, ultimately driving the molecular mechanisms that underlie disease pathogenesis.

In summary, the development of a suite of chemigenetic biosensors with far-red emission and high dynamic range extends the capacity of imaging signal dynamics from cellular levels to intact tissues, providing a versatile and powerful toolkit with multiplexed capability across spectral windows. These advances enable the detection of delicate signaling dynamics in native contexts, thereby accelerating our understanding of disease etiology and establishing a framework for the development of signaling-targeted therapeutic strategies.

## Methods

### Biosensor construction

To construct RHt-AKAR, HaloTag was PCR-amplified from pcDNA5/FRT/TO_H2B_HaloTag_T2A_EGFP^72^ using the forward primer 5’-CAGATCGCTGAGATAGGTGCC-3’ and the reverse primer 5’-ACCGGAAATCTCCAGAGTAGACAG-3’. In parallel, the AKAR4^7^ scaffold (Cerulean, FHA1, and PKA substrate sequence) was PCR-amplified using the forward primer 5’-CTGTCTACTCTGGAGATTTCCGGTTAAGAATTCTGCAGATATCCAGCACAG-3’ and the reverse primer 5’-GCCAGTACCGATTTCGGATCCCATGAGCTCGCTGCCG-3’. The two fragments were combined using Gibson assembly. The resulting plasmid was re-linearized by PCR-amplification using the forward primer 5’-ACAAGAGATCTATGCATAAGTTTTCTCAAGAAC-3’ and the reverse primer 5’-CAGATCGCTGAGATAGGTGCC-3’. sTagRFP was PCR-amplified from pcDNA3-sTagRFP^31^ using the forward primer 5’-ATGAGCGAGCTGATTAAGGAGAAC-3’ and the reverse primer 5’-CTTGTGCCCCAGTTTGCTAG-3’. These two fragments were assembled by Gibson to yield RHt-AKAR. YHt-AKAR was generated by PCR amplification of mCitrine from ERT2-PhoCI-Inactive-PKI-PhoCI-ERT2 and pCre-mCitrine^73^ using the forward primer 5’-TTGCGGCCGCCACC<u>GGATCC</u>ATGGTGAGCAAGGGCGAG-3’ and the reverse primer 5’-GAAAACTTATGCAT<u>AGATCT</u>CTTGTACAGCTCGTCCATG-3’ (overlaps corresponding to BamHI and BglII sites are underlined). Each donor variants of YHt-AKAR and RHt-AKAR were generated with a similar cloning strategy. The PCR product was assembled by Gibson into BamHI/BglII-digested RHt-AKAR backbone. Vector backbones containing PKC substrate, Epac1 (for cAMP), Pseudoligand (for Ras), and Src substrate sequences were similarly used to generate RHt-CKAR, YHt-CKAR, RHt-ICUE, YHt-ICUE, YHt-RasAR, and YHt-SrcAR. Ht-AKARs EV linker variants were generated as follows. Constructs containing 258 aa EV linkers were generated by ligating a KpnI/BspEI-digested 258 aa EV linker fragment from Booster-PKA^30^ into recipient backbones digested with the same restriction enzymes. For other linker lengths, EV linker segments were PCR-amplified from AKAR3-EV^26^ using the forward primer 5’-GCAGAACAAAGTTGATCGC<u>GGTACC</u>AGTGCTGGTGGTAGTGCTGGTGGTAGTGCTGGTGGTAGTGCTGGTGGT AGTGCTGGTGGTTCCGGCAGTGCTGGTGGTAGTGCTGGTGGTAGTACCAGTGCTGGTGGTAGTGCTG-3’ and the reverse primer 5’-CGGAAGGCCTGGAAGGTCTC<u>TCCGGA</u>ACCACCAGC-3’ (overlaps corresponding to KpnI and BspEI sites are underlined). The resulting PCR products were assembled by random Gibson assembly into KpnI/BspEI-digested backbones. Multiple colonies were tested to obtain constructs spanning different EV linker lengths. All constructs were verified by sequencing.

### Cell culture and transfection

HeLa cells were cultured in Dulbecco’s modified Eagle medium (DMEM, Gibco) containing 1 g L^−1^ glucose and supplemented with 10% (v/v) fetal bovine serum (FBS, Sigma-Aldrich) and 1% (v/v) penicillin-streptomycin (Sigma-Aldrich). HEK293T and COS-7 cells were cultured in DMEM (Gibco) containing 4.5 g L^−1^ glucose, 10% (v/v) FBS (Sigma) and 1% (v/v) penicillin-streptomycin (Sigma-Aldrich). All cells were maintained in a 37 °C incubator with a humidified 5% CO_2_ atmosphere. All cell lines were determined to be free of mycoplasma contamination based on weekly DNA staining. For fluorescence imaging experiments, cells were seeded onto sterile quartered 35-mm glass-bottomed dishes (Cellvis) and grown to 50%-70% confluence for transient transfection. For HEK293T cells, dishes were poly-D-lysine (ThermoFisher) coated before seeding. Transient transfection of HeLa cells were performed using Lipofectamine 2000 (Invitrogen) according to the manufacturer’s recommendations. The solutions were incubated for 5 min at room temperature, then mixed and incubated for 10 min. The prepared DNA-Lipofectamine complex was added to one of the wells and incubated for 6 h, after which medium was changed to fresh medium. HEK293T and COS-7 cells were transfected using Polyjet (SignaGen) with the same processes as Lipofectamine 2000 without a fresh medium change. Prior to transfection, the COS-7 cells were washed with Dulbecco phosphate-buffered saline (DPBS, Gibco), and medium was changed to a serum-free DMEM to serum-starve the cells for 24 h during transfection prior to imaging. All cells were grown for an additional 20-24 h before imaging.

### Labeling and preparation for widefield live-cell imaging

Cells were labeled with 500 nM JF_669_ (L. Lavis Lab at Janelia Research Campus) for 4 h at 37 °C in the appropriate pre-warmed cell culture medium. Cells were then washed twice and incubated in Hank’s balanced salt solution (HBSS, Gibco) before imaging. For PKA and cAMP imaging in HeLa and HEK293T or Ras and Src imaging in COS-7, cells were imaged immediately. For PKC imaging in HeLa, cells were additionally incubated in HBSS for 15 min at 37 °C prior to imaging.

### Live-cell epifluorescence microscopy

Cells were imaged in the dark at 37°C on a Zeiss AxioObserver Z1 microscope (Carl Zeiss) equipped with a 40x/1.3 NA objective, a Photometrics Evolve 512 EMCCD (Photometrics) controlled by METAFLUOR 7.10 software (Molecular Devices). An OD filter of OD = 0.6 was used unless otherwise stated. Forskolin (Fsk, Calbiochem, #344281), 3-isobutyl-1methylxanthine (IBMX, Sigma-Aldrich, #I5879), H89 (Cayman Chem, #10010556), phorbol 12-myristate 13-acetate (PMA, LC Laboratories, #P1680), Gö6983 (Sigma, #G1918), epidermal growth factor (EGF, Sigma-Aldrich, #E9644), isoproterenol (iso, Sigma-Aldrich, #I5627), propranolol (Prop, Sigma, #P0884), carbachol (Thermo Scientific, #AAL0667403), pertussis toxin (PTX, Sigma, #516561), FR900359 (Cayman Chemical, #33666) were added as indicated.

Dual emission ratio imaging was performed using an ET495/10x excitation filter, 515dcxr dichroic mirror, and HQ535/25m (yellow) and ET700/75m (far-red) emission filters for YHt-sensors (Y/FR) and an ET555/25x excitation filter, ZT568rdc dichroic mirror, and ET650/100m (red) and ET700/75m (far-red) emission filters for RHt-sensors (R/FR). Dual GFP excitation ratio imaging of ExRai-CKAR was performed using ET405/40x and HQ480/30x excitation filters, a 505dcxr dichroic mirror, and an HQ535/45m emission filter. Exposure times were 100 ms (far-red channel) and 500 ms (all other channels). Images were acquired every 30 s, except for multiplexing of RHt-AKAR with YHt-CKAR, which was every 20 s.

Raw fluorescence images were corrected by subtracting the background fluorescence intensity of a cell-free region from the emission intensities of biosensor-expressing cells. The corresponding emission ratios for chemigenetic biosensors and excitation ratios for ExRai-CKAR were then calculated at each time point. The time-course data were normalized by dividing the ratio at each time point by the basal value at time zero (*R/R*_0_), which was defined as the last time point immediately preceding drug treatment. Microsoft Excel 365 was used to process data. Images were processed with METAFLUOR. Graphs were plotted using GraphPad Prism 10 (GraphPad Software).

### Two-photon FLIM (2p-FLIM) imaging

2p-FLIM imaging was performed on a custom-built microscope equipped with a mode-locked laser (Spectra-Physics, Insight X3 operating at 80 MHz). Emitted photons were detected by fast photomultiplier tubes (PMTs, Hamamatsu, H10770PB-40). For cellular resolution imaging, a 60× [Olympus, numerical aperture (NA) 1.1] objectives were used, respectively. Image acquisition used custom-written software ScanImage^74^ in MATLAB 2012b. FLIM was performed as described previously^63,75^. 1100 nm and 1050 nm were used as the excitation wavelength at 2.5 mW for HEK293T cells and brain slices, respectively. A 594LP dichroic (DI02-R594-25X36, Semrock) and a 600/50 nm band-pass filter (ET600/50m, Chroma) were used to collect the emission light of donor fluorescence from RHt-AKAR. During the experiments, RHt-AKAR lifetime was recorded for over 5 min as a baseline. Isoproterenol (1 μM) and Fsk (50 μM) were added directly to the perfusion reservoir.

#### In utero electroporation of RHt-AKAR

The mouse husbandry and surgery were performed following the approved protocols by the Institutional Animal Care and Use Committee of Washington University in St. Louis and in accordance with National Institutes of Health guidelines. In utero electroporation was performed on an E15 pregnant CD-1 mouse under 2% isoflurane anesthesia to express RHt-AKAR in the left somatosensory cortex of the pups. The embryo head was then positioned with a tweezers fitted with round plate electrodes (0.5 mm diameter), and electrical pulses were applied at a rate of five times per second (50 V, 50 ms), with the cathode on the right cortex and the anode at the left cortex (CUY21 electroporator, NEPA GENE, Japan). For each pup, 1 μg of pBS-β-actin-Cre (gift from S. Dymecki Lab) and 2 μg of pAAV-CAG-DIO-RHt-AKAR were electroporated. Warm PBS was intermittently dripped onto the embryos. The uterus was then returned back into the pregnant mother, and the muscle and the skin were sutured separately. Pups were housed with the mother until use.

#### Imaging in HEK293T cells

Cells were seeded on coverslips in 24-well tissue culture plates and then transfected the following day with plasmids (0.3 μg pAAV-CAG-DIO-RHt-AKAR, 0.2 μg β-actin-Cre per well (gift from S. Dymecki Lab)) with Lipofectamine 2000 (Invitrogen). Two days after transfection, 2p-FLIM was performed. Coverslips were incubated with 50 nM JF_669_ (gift from L. Lavis Lab at Janelia Research Campus) for at least 30 mins prior to imaging. During imaging, coverslips were perfused with HEPES buffer (138 mM NaCl, 1.5 mM KCl, 1.2 mM MgCl_2_, 2.5 mM CaCl_2_, 10 mM HEPES, 10 mM glucose, pH with NaOH to 7.35) at 32°C and a flow rate of 2 mL/min.

#### Preparation and imaging of acute brain slices

P16-18 mice were anesthetized with isoflurane before being sacrificed. Their brains were then rapidly dissected out. Acute horizontal sections were sliced from the hippocampus with a Leica VT1000S vibratome (Leica Instruments) in cold sucrose cutting solution (87 mM NaCl, 25 mM NaHCO_3_, 1.25 mM NaH_2_PO_4_, 2.5 mM KCl, 1 mM MgCl_2_•6H_2_O, 25 mM glucose, 75 mM sucrose). Slices were 300 μm thick. After sectioning, slices were transferred to artificial cerebrospinal fluid (ACSF; 127 mM NaCl, 25 mM NaHCO_3_, 1.25 mM NaH_2_PO_4_, 2.5 mM KCl, 1 mM MgCl_2_•6H_2_O, 2 mM CaCl_2_) at 34°C for 20 min, and then incubated in dye solution containing 1 µM JF_669_ in ACSF at room temperature for at least 30 mins. The brain slices were perfused with ACSF with a 2-4ml/min flow rate throughout the experiments. The cutting solution, ACSF, and dye solution were bubbled with carbogen (5% CO_2_, 95% O_2_).

#### Two-photon FLIM data analysis

The cytoplasm of the neurons was analyzed. The decay constant for donor fluorophore in free form, denoted as τ_1_, was determined as 2.2 ns with a single exponential equation to fit the lifetime histogram of the cytoplasm of HEK293T cells transfected with donor only. The decay constant for donor in FRET form, denoted as τ_2_, was best fitted to be 0.7 ns through double exponential fitting of the lifetime response of RHt-AKAR expressed in HEK293T cells with τ_1_ = 2.2 ns. The same τ_1_ and τ_2_ values were used for lifetime fitting for brain slice experiments. The lifetime change in baseline was calculated as the difference between the average of the first five data points and the average of the last five data points during the baseline recording. The lifetime change after drug application was calculated as the difference between the average of the last five data points during drug perfusion and the average of the last five data points of the baseline recording. Microsoft Excel 365 was used to process data. Images were processed with ImageJ/Fiji.

### Statistics and reproducibility

Statistical analyses were performed using GraphPad Prism 10 (GraphPad Software). Statistical significance was set at *P* < 0.05. All experiments were independently repeated at least three times with similar results. Representative microscopy images and time-courses are shown. Cells to be analyzed were chosen on the basis of the proper localization of biosensor fluorescence, reflecting correct cell functionality in response to agonist stimulation, and cells displaying clearly unusual behavior (such as apoptotic blebbing) were excluded from analysis. Cells were randomly seeded into experimental groups after resuspension for all cell-based data. Fields of view were randomly chosen within imaging dishes.

### Data Availability

Biosensor constructs generated in this study will be made available through Addgene (www.addgene.org/Jin_Zhang). Source data are provided with the manuscript.

## Acknowledgements

The authors are grateful to Luke Lavis and Jonathan Grimm (Janelia Research Campus) for generously providing JF dyes. We thank Asuka Inoue for providing the GNAS knockout HEK293A, Wei Lin for help with imaging experiments, Yingqi Zhu, Nick D. Bergkamp, and Jinfan Zhang for assistance with cloning and providing plasmids, and Qiang Ni for support in material acquisition and tissue culture. This work is supported by the National Institutes of Health (grants R35 CA197622, R01 DK073368, R01 DE030497, R01 CA262815, and RF1 MH126707 to J.Z; R01 NS119821 to Y.C; T32 NS121881 to Z.B.R), by the Swiss National Science Foundation (SNSF) (grants P2ELP3_199834 and P500PN_214234 to M.S.F), as well as by the Kavli Foundation (grants LS-2024-GR-01-2898 to Y.C).

## Author Contributions

X.H., M.S.F., and J.Z. conceived the project. X.H., M.S.F., Y.C., and J.Z. designed the experiments. X.H., D.R.Z., and M.S.F. generated biosensors. X.H. performed live-cell imaging in HeLa, HEK293T, and COS-7 cells with assistance from M.S.F. and D.R.Z. Y.H. performed FLIM imaging in brain slices. Z.B.R performed FLIM imaging in HEK293T. X.H., Y.H., Z.B.R., and D.R.Z. analyzed data. J.Z. coordinated the study and provided guidance. X.H., M.S.F., S.M., and J.Z. wrote the paper. All authors discussed the results and approved the final version of the manuscript.

