## Supplementary Information for "Highly Sensitive Chemigenetic FRET-Based Kinase Biosensors"

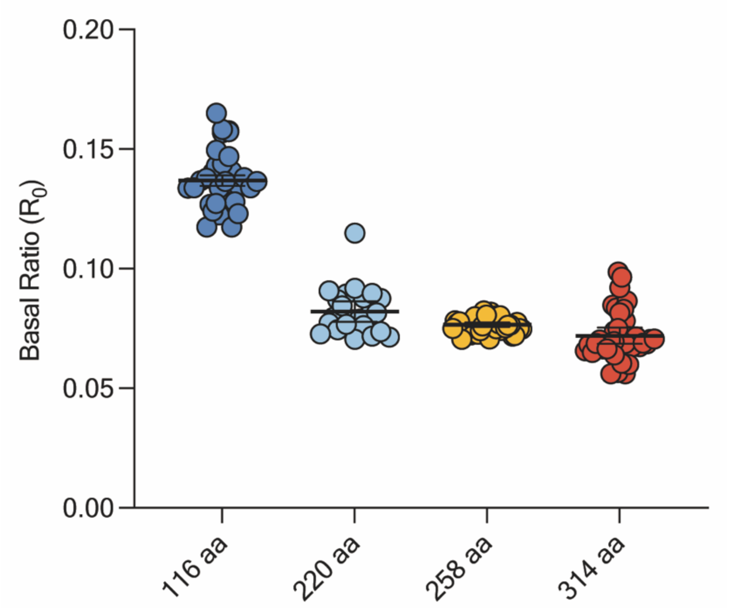


**Supplementary Figure S1:** Quantification of basal emission ratio ($R_0$) of YHt-AKAR EV linker variants. n = 32 (116 aa), n = 23 (220 aa), n = 43 (258 aa), and n = 38 cells (314 aa). Individual data points are pooled from three independent experiments. Error bars indicate mean ± SEM.

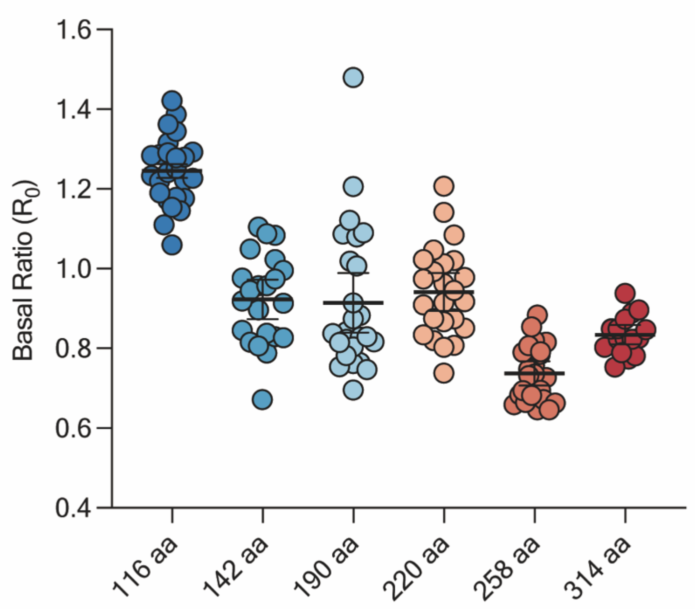


**Supplementary Figure S2:** Quantification of basal emission ratio ($R_0$) for RHt-AKAR EV linker variants. n = 25 (116 aa), n = 22 (142 aa), n = 25 (190 aa), n = 24 (220 aa), n = 23 (258 aa), and n = 17 cells (314 aa). Individual data points are pooled from three independent replicates. Error bars indicate mean ± SEM.

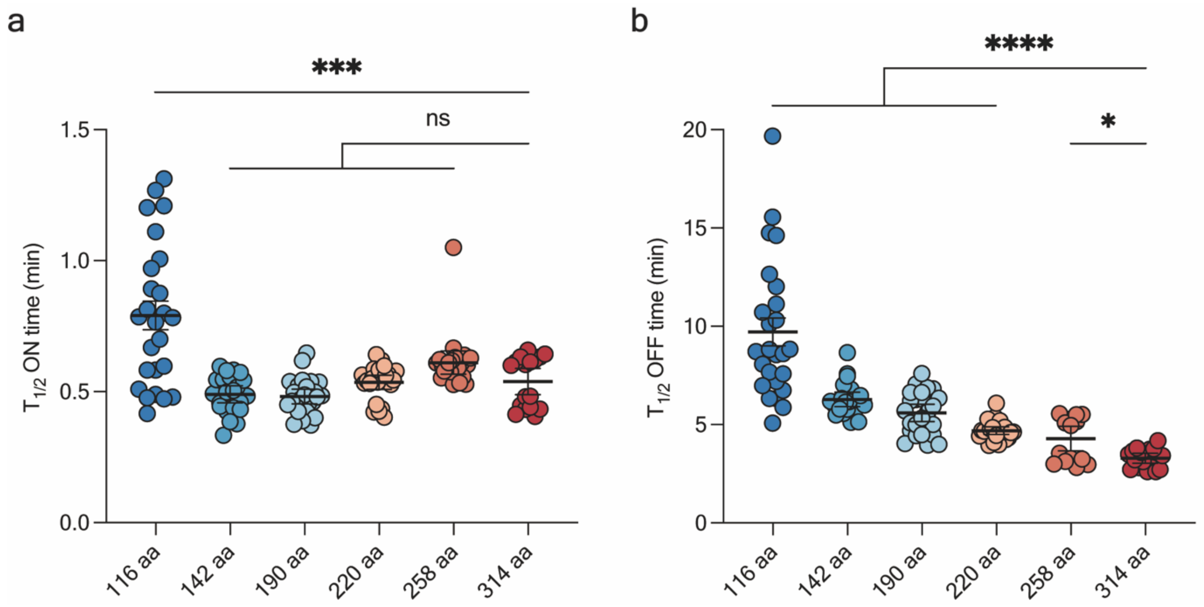


**Supplementary Figure S3:** Quantification of response kinetics of RHt-AKARs EV linker variants. **a,** Time to half-maximal response ($T_{1/2}$ ON) after 50 µM Fsk and 100 µM IBMX addition. n = 25 (116 aa), n = 22 (142 aa), n = 25 (190 aa), n = 24 (220 aa), n = 23 (258 aa), and n = 17 cells (314 aa). **b,** Time to half-maximal inhibition ($T_{1/2}$ OFF) after 20 µM H89 addition. n = 25 (116 aa), n = 22 (142 aa), n = 25 (190 aa), n = 24 (220 aa), n = 15 (258 aa), and n = 17 cells (314 aa). Individual data points are pooled from three independent experiments. Error bars indicate mean ± SEM. Data were analyzed using one-way ANOVA followed by Dunnett's multiple-comparison test, ****$P < 0.0001$, ***$P = 0.0009$, *$P = 0.0243$, ns = nonsignificant different.

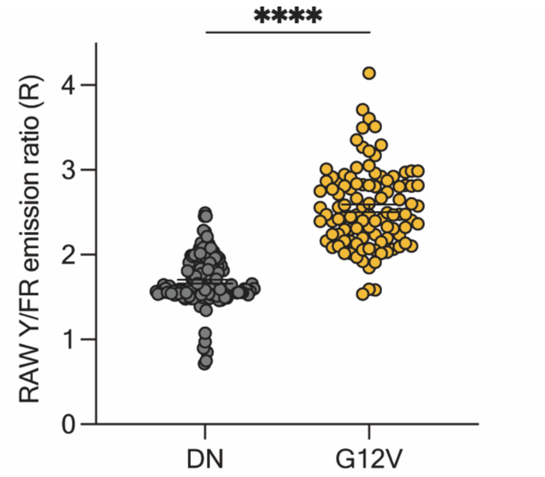


**Supplementary Figure S4:** YHt-RasAR characterization. Quantification of YHt-RasAR emission ratio in COS-7 cells co-expressing either dominant-negative HRas S17N (DN) (n = 149) or constitutively active HRas G12V (G12V) (n = 104). Individual data points are pooled from two independent experiments. Error bars indicate mean ± SEM. Data analyzed using unpaired, two-tailed Student's t-test, ****$P < 0.0001$.

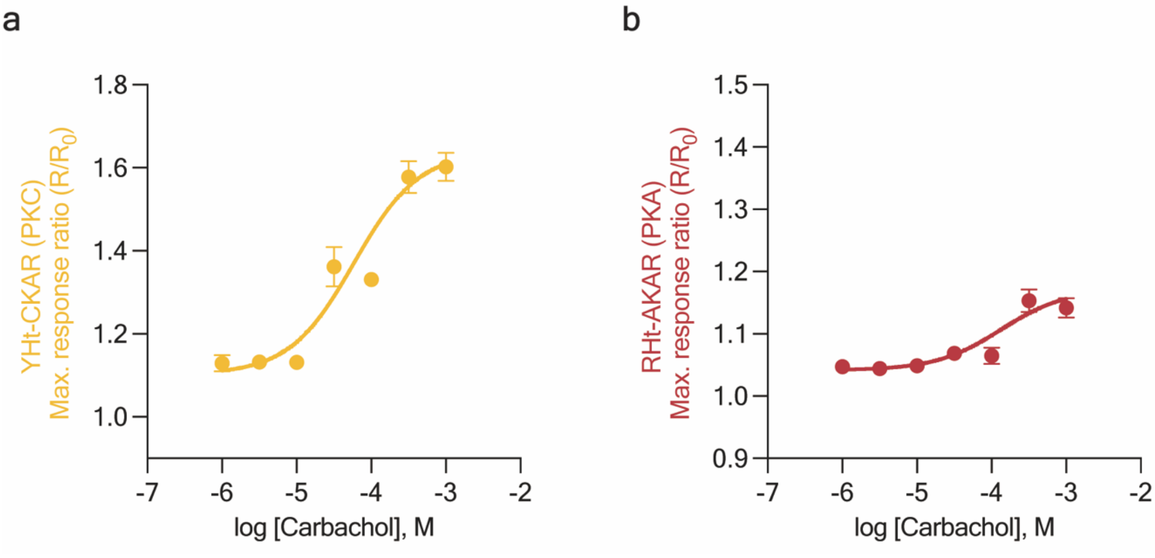


**Supplementary Figure S5:** Concentration response curves showing **(a)** YHt-CKAR and **(b)** RHt-AKAR responses upon stimulation with different concentrations of carbachol (1 µM, 3 µM, 10 µM, 30 µM, 100 µM, 300 µM, and 1 mM) in M2R-overexpressing HEK293T cells. Data points represent mean ± SEM from two independent experiments along with mean and SEM. From lowest to highest concentration: n = 25, 29, 31, 26, 13, 31, 21 cells.

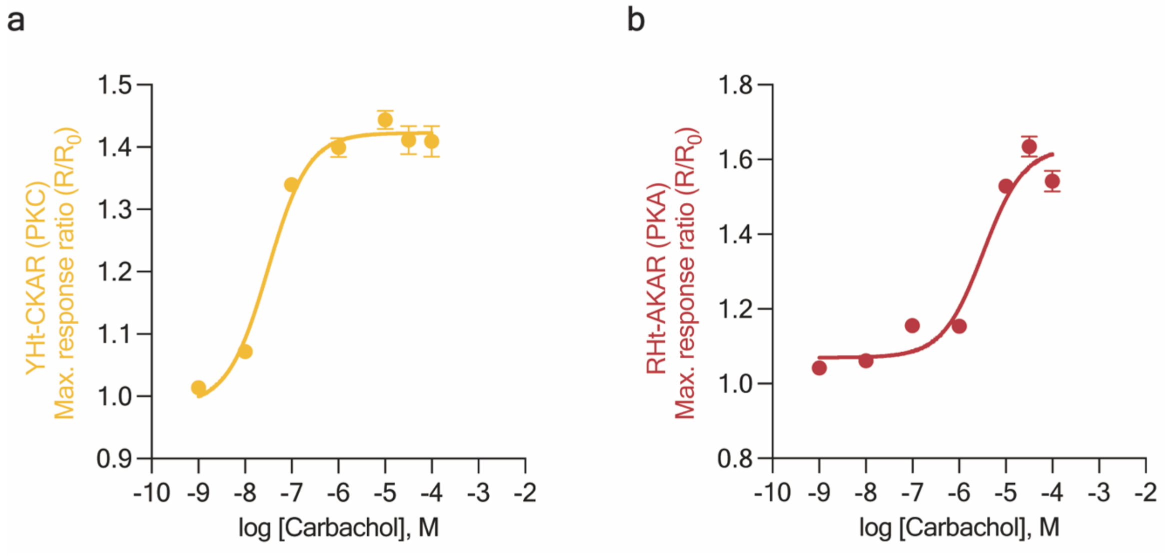


**Supplementary Figure S6:** Concentration response curves showing **(a)** YHt-CKAR and **(b)** RHt-AKAR responses upon stimulation with different concentrations of carbachol (1 nM, 10 nM, 100 nM, 1 µM, 10 µM, 30 µM, and 100 µM) in M3R-overexpressing HEK293T cells. Data points represent mean ± SEM from two independent experiments. From lowest to highest concentration: n = 20, 25, 25, 32, 26, 22, 21 cells.

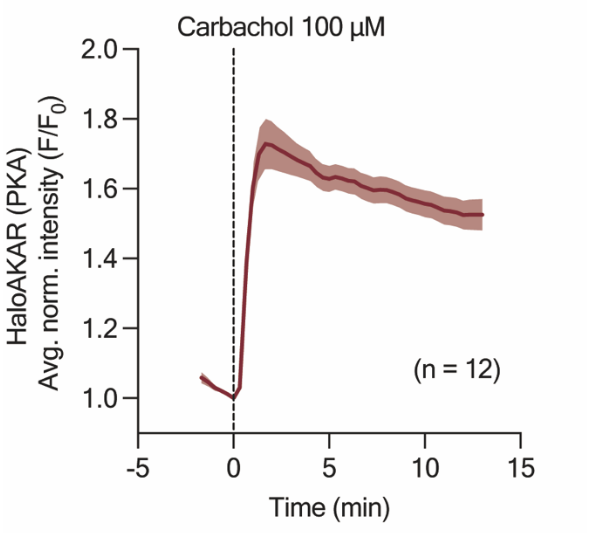


**Supplementary Figure S7:** Endogenous M3R-mediated PKA activation reported by HaloAKAR. Representative average time-course showing HaloAKAR response in HEK293A cells treated with 100 µM carbachol. Data are representative of two independent experiments. Dashed line indicates drug addition, solid lines indicate mean response, and shaded areas depict SEM.

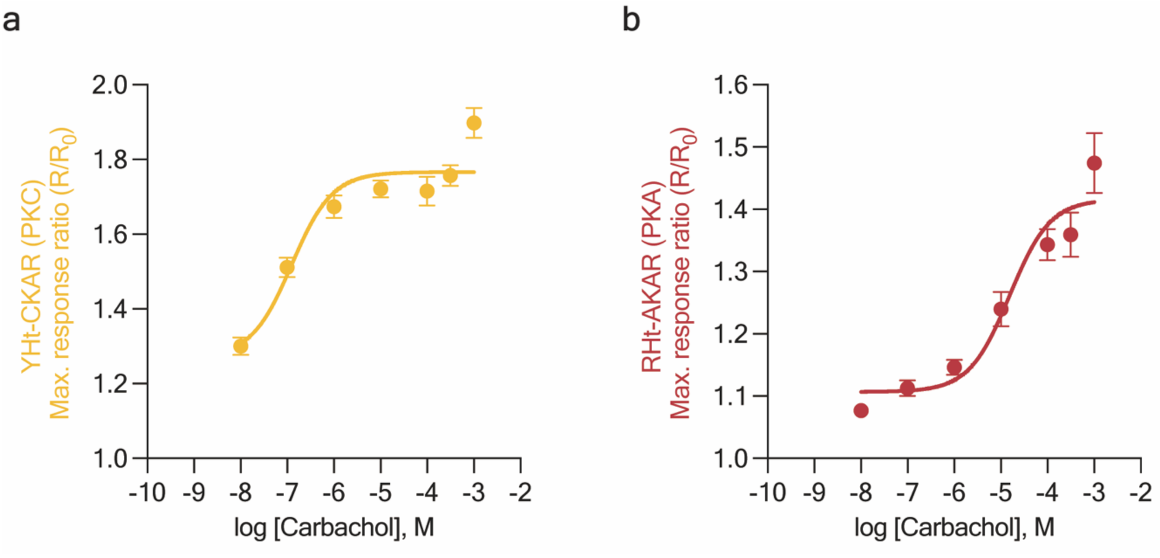


**Supplementary Figure S8:** Concentration response curves showing **(a)** YHt-CKAR and **(b)** RHt-AKAR responses upon stimulation with different concentrations of carbachol (10 nM, 100 nM, 1 µM, 10 µM, 100 µM, 300 µM, and 1 mM) in M5R-overexpressing HEK293T cells. Data points represent mean ± SEM from two independent experiments. From lowest to highest concentration: n = 25, 28, 29, 31, 25, 27, 26 cells.

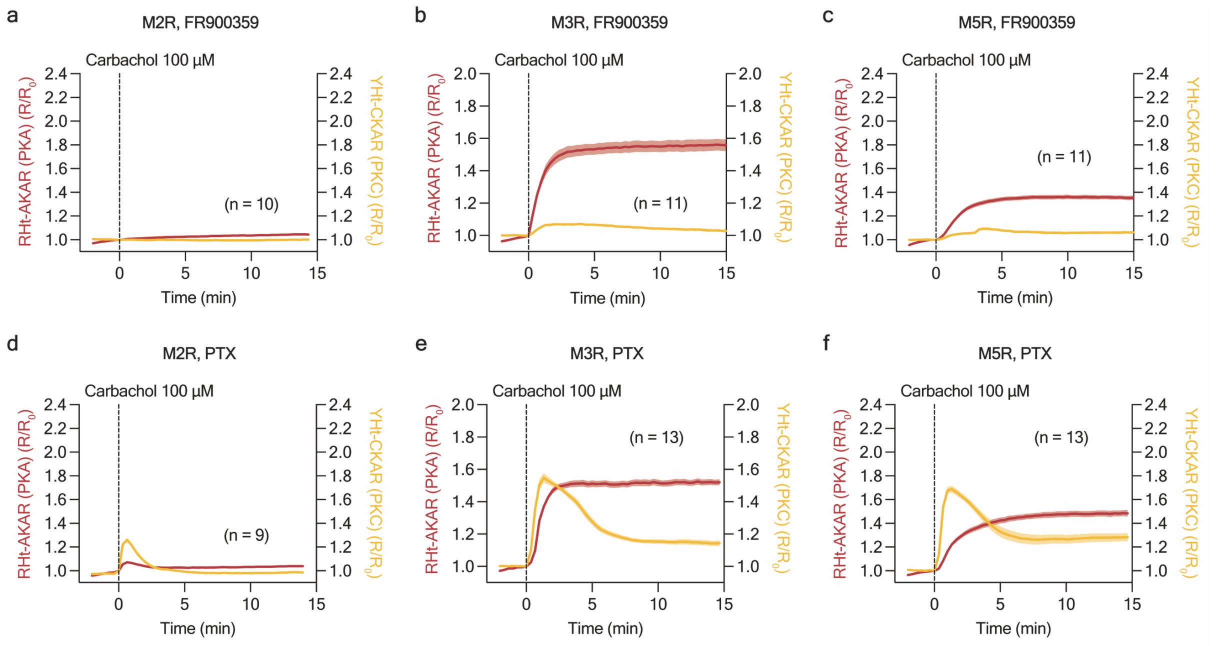


**Supplementary Figure S9:** Multiplexed activity imaging of PKA and PKC activity downstream of different mAChR isoforms in HEK293T cells. **a-f,** Representative average time-courses of RHt-AKAR (red) and YHt-CKAR (yellow) responses to 100 µM carbachol stimulation in HEK293T cells co-expressing M2R **(a)**, M3R **(b)**, or M5R **(c)** and pretreated with 100 nM FR900359 for 30 min, or co-expressing M2R **(d)**, M3R **(e)**, or M5R **(f)** and pretreated with 100 ng/mL PTX (6 h). Time-courses are representatives of two independent experiments. Dashed lines indicate drug addition, solid lines indicate mean responses, and shaded areas depict SEM.

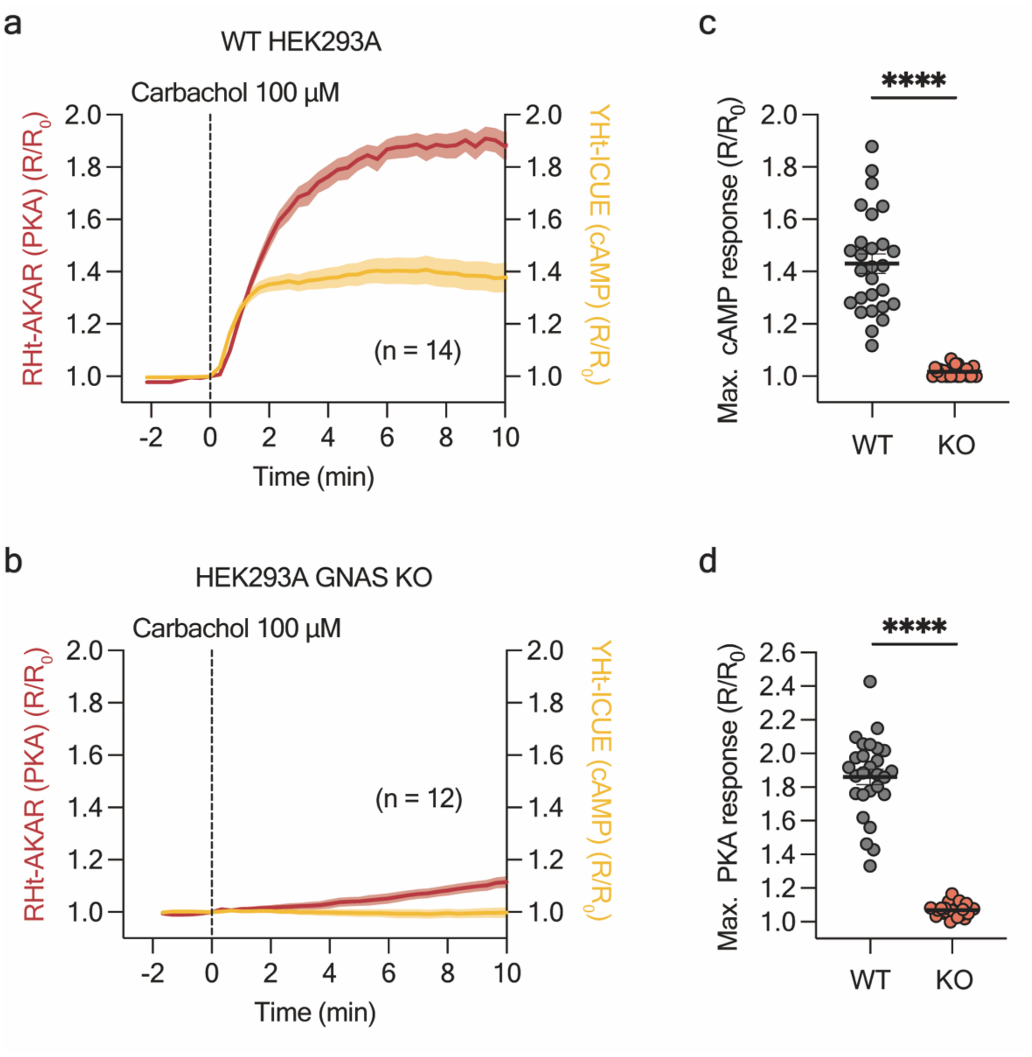


**Supplementary Figure S10:** Multiplexed activity imaging of M3R-induced PKA and cAMP dynamic in HEK293A cells. **a-b,** Representative average time-courses of RHt-AKAR (red) and YHt-ICUE (yellow) responses to 100 µM carbachol stimulation inHEK293A **(a)** or HEK293A GNAS KO cells **(b)** co-expressing M3R. Time-courses are representatives of two independent experiments. Dashed lines indicate drug addition, solid lines indicate mean responses, and shaded areas depict SEM. **c-d,** Quantification maximum cAMP **(c)** or PKA **(d)** responses 5 min post carbachol stimulation in HEK293A (parental, n = 27) or HEK293A GNAS KO (n = 24) cells overexpressing M3R. Data points are pooled from two independent experiments. Error bars indicate mean ± SEM. Data were analyzed using unpaired, two-tailed Student's t-tests, ****$P < 0.0001$.
